# Single-cell Rapid Capture Hybridization sequencing (scRaCH-seq) to reliably detect isoform usage and coding mutations in targeted genes at a single-cell level

**DOI:** 10.1101/2024.01.30.577942

**Authors:** Hongke Peng, Jafar S. Jabbari, Luyi Tian, Chong Chyn Chua, Natasha S. Anstee, Noorul Amin, Andrew H. Wei, Nadia M. Davidson, Andrew W. Roberts, David C. S. Huang, Matthew E. Ritchie, Rachel Thijssen

**Author notes:** Correspondence to R. Thijssen; at the Department of Hematology, Amsterdam UMC, De Boelelaan 117, Amsterdam, The Netherlands. Joint senior author (Matthew E. Ritchie, Rachel Thijssen).

## Abstract

Single-cell long-read sequencing has transformed our understanding of isoform usage and the mutation heterogeneity between cells. Despite unbiased in-depth analysis, the low sequencing throughput often results in insufficient read coverage thereby limiting our ability to perform mutation calling for specific genes. Here, we developed a **s**ingle-**c**ell **Ra**pid Capture **H**ybridization **seq**uencing (scRaCH-seq) method that demonstrated high specificity and efficiency in capturing targeted transcripts using long-read sequencing, allowing an in-depth analysis of mutation status and transcript usage for genes of interest. The method includes creating a probe panel for transcript capture, using barcoded primers for pooling and efficient sequencing via Oxford Nanopore Technologies platforms. scRaCH-seq is applicable to stored and indexed single-cell cDNA which allows analysis to be combined with existing short-read RNA-seq datasets. In our investigation of BTK and SF3B1 genes in samples from patients with chronic lymphocytic leukaemia (CLL), we detected SF3B1 isoforms and mutations with high sensitivity. Integration with short-read scRNA-seq data revealed significant gene expression differences in SF3B1-mutated CLL cells, though it did not impact the sensitivity of the anti-cancer drug venetoclax. scRaCH-seq’s capability to study long-read transcripts of multiple genes makes it a powerful tool for single-cell genomics.

## Introduction

Single-cell sequencing technologies have revolutionized our understanding of cell state and function(1,2). These approaches enable the comprehensive characterization of individual cells, revealing cancer biology and tumour heterogeneity(3-9). While single-cell RNA sequencing (scRNA-seq) has been widely used for transcriptomic profiling of individual cells, it has limitations in calling mutations and quantifying isoform usage due to the 5’ and 3’ bias induced by the fragmentation step in the library preparation protocol and the widespread use of short-read technology for sequencing samples. Consequently, linking the single-cell transcriptome to the mutation status of cancer cells becomes a challenge. Understanding the clonal and non-clonal mechanisms of selection and adaptation in response to therapeutic pressure is of importance. Moreover, the influence of isoform usage on cell states further underscores the need for comprehensive methodologies. In response to these challenges, alternative methods have emerged to link the transcriptome profile to isoform usage, mutations, and translocations in full-length transcripts at a single-cell level(10).

In recent years, several high-throughput methods have been developed to enable single-cell long-read sequencing. These approaches typically involve barcoding of cDNA using existing methods such as 10x Genomics or Drop-seq and sequencing the indexed full-length cDNA on platforms such as Pacific Biosciences (PacBio) or Oxford Nanopore Technologies (ONT)(11,12). While these unbiased single-cell long-read sequencing methods can detect a larger number of isoforms at a single-cell level, the lower overall sequencing level per cell reduces the ability to accurately quantify isoform usage and mutation calling. To overcome this challenge, other methods were developed to target genes of interest, but these methods often require primer spike-in with the 10x Genomics protocol and primer panel design optimization(13,14). This limits the ability to target multiple genes of interest or to perform long-read sequencing on already amplified single-cell indexed cDNA.

The growing interest in high throughput single-cell multi-omic methods to enhance our understanding of cancer complexity led us to develop a method that can flexibly link genotypes in single cells to pre-existing short-read transcriptome data generated by various 10x Genomics protocols. CITE-seq combines transcriptome with cell surface protein expression and 10x Genomics multiome combines transcriptome with chromatin accessibility data(15). Our single-cell Rapid Capture Hybridization sequencing (scRaCH-seq) method enables the capture of multiple transcripts from pre-indexed and stored cDNA independent of the 10x Genomics kit used. This approach uses biotinylated probes to target genes of interest and Streptavidin beads for on-target transcript enrichment. It was first detailed in Thijssen *et al*. (Blood 2022), to capture the transcriptional landscape of 17 BCL2 family genes in patients with chronic lymphocytic leukaemia (CLL)(16). That study investigated the resistance mechanisms observed in chronic lymphocytic leukaemia cells after treatment with the BCL2 inhibitor venetoclax treatment. Using scRaCH-seq, we successfully identified the BCL2 G101V mutation, a novel PMAIP1 transcript and a unique BAX isoform across different venetoclax-relapsed samples at a single-cell level. Despite the first application of scRaCH-seq to that research, a detailed method description, data analysis and method evaluation has not been published.

Here, we provide a comprehensive description of the scRaCH-seq method in which we target the 2 genes SF3B1 and BTK as illustrative examples. Among the CLL patients treated with venetoclax, SF3B1 mutation was detected in 5 samples by whole exome sequencing (WES)(16) which was used as a ground truth to validate the scRaCH-seq data. A step-by-step guide to probe panel design was created to facilitate the application of scRaCH-seq together with a tailored analysis pipeline that extends the FLAMES (**F**ull-**L**ength **A**nalysis of **M**utations and **S**plicing) R package (17) and incorporates additional read integrity checks and steps to remove background noise. The performance of the scRaCH-seq method was carefully evaluated, focusing on key aspects including probe capture efficiency, artefact diagnostics and mutation/deletion calling.

## Materials and methods

### Samples

The full-length cDNA of 18 peripheral blood samples from chronic lymphocytic leukaemia (CLL) patients and 3 healthy donors (HD) were used(16). 10x Genomics was performed using the Chromium Next GEM (Gel Bead-In Emulsions) Single Cell V(D)J (v1.1, 10x Genomics, cat#PN-1000165) according to the manufacturer’s instructions. The indexed full-length surplus cDNA(16) was stored and used as input for this scRaCH-seq experiment (**Figure 1**).

**Figure 1.**
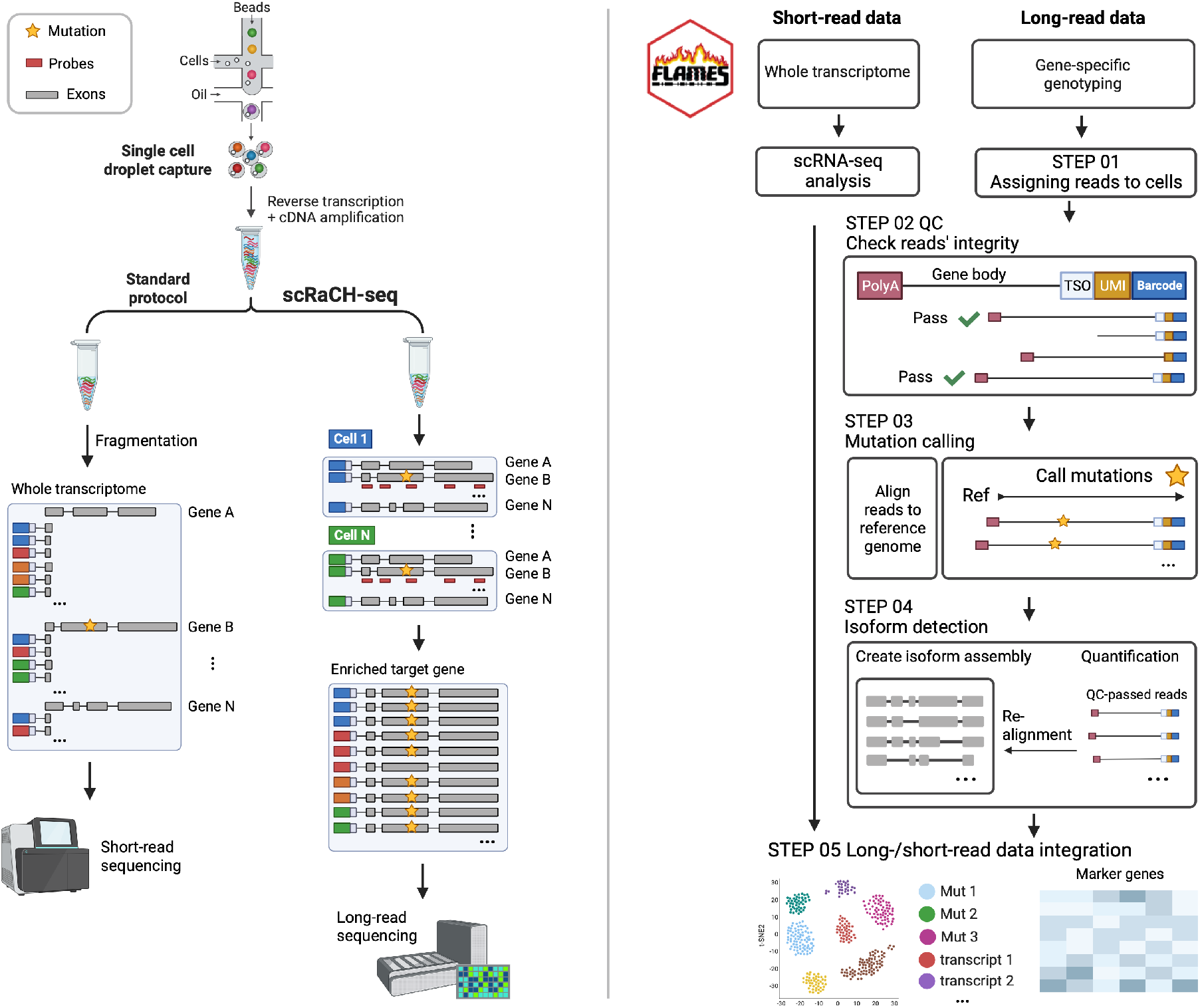
Schematic of scRaCH-seq approach and FLAMES pipeline for single-cell long-read data analysis. scRaCH-seq can be incorporated in the standard high-throughput single-cell RNA-seq experiments (left side). After single-cell isolation, mRNA is converted into cDNA and barcoded. Only some of this amplified indexed cDNA is used for short-read library preparation and Illumina sequencing. The surplus cDNA will be stored and can be used for scRaCH-seq. For scRaCH-seq, a probe panel will be designed for genes of interest. The biotinylated probes will be hybridized with amplified cDNA overnight. The probes and target genes will be captured with Streptavidin beads, washed, and amplified. The enriched target long-read transcripts will be sequenced on the Nanopore platform. With the FLAMES pipeline, Fastq files were demultiplexed by cross-referencing the cell barcodes identified in scRNA-seq data (STEP 01). Next, the demultiplexed reads will undergo an integrity check and the reads that possess a UMI, a TSO sequence, and poly-A tails are retained (STEP 02). Reads were aligned to the GRCh38 reference genome to construct a transcript assembly. Concurrently, the piled-up reads were compared against GRCh38 to identify base alterations and deletions, generating a comprehensive mutation/deletion matrix (STEP 03). The reads will be realigned to the transcript assembly for quantification (STEP 04). Single-cell short-read gene expression data was used to cluster the cells, based on the conventional analysis pipeline for scRNA-seq. The transcript usage and mutation/deletion information were then aligned with the single-cell gene expression data. The bridge connecting these datasets was the shared cell barcodes, ensuring the gene expression profile at a transcript level (STEP 05).

### Probe panel design

Probes were designed to capture the indexed cDNA of target genes. Each probe was 120 base pairs (bp) in length, ensuring robust coverage of the targeted regions. First, a fasta file was generated for all the isoforms of the genes of interest. The fasta file containing all Ensembl-annotated exons from all the isoforms was compiled and this was achieved by extracting the corresponding sequences from the human genome GRCh38. This fasta file served as the foundation for probe design. Next, exon sequences shorter than 120 bp were merged with the preceding and subsequent exons to create concatenations exceeding 120 bp, resulting in an updated fasta file with all reads over 120 bp. Using the custom fasta file, a probe panel was generated to cover each base in the input, employing https://sg.idtdna.com/pages/tools/xgen-hyb-panel-design-tool. For probe selection, a strategy was implemented to cover every 1000 bp of an exon sequence with a single probe (**Supplementary Figure 1**). Probes were selected based on their GC content (GC%) to ensure consistent efficiency during their hybridization with cDNA. The average GC% of the redundant probe panel was used as a baseline. The probe with GC% closest to the average GC% was chosen from the group of probes covering a thousand-base region. For concatenated exons, probes fully covering the entire sequence of the original short exon were selected (**Supplementary Figure 1)**. Duplicated probes in the selected panel were removed to finalize the probe panel. A pre-fixed R script detailing the code for the probe panel design process as well as a step-by-step instruction is available at the following GitHub repository: https://github.com/HongkePn/RaCHseq_Probe_Design.

### Single-cell Rapid Capture Hybridization sequencing (scRaCH-seq)

#### Amplification of cDNA

The surplus 10x indexed cDNA was amplified to increase the cDNA yield for the hybridization step. To obtain (1.5-2 μg) cDNA, 2-10ng of cDNA was used as input in 5x50μl reactions. For a 50μl reaction using 10× Genomics 3’ or 5’ cDNA, a mixture containing 10μl 5x PrimeSTAR buffer (Takara, #R050B), 4μl dNTP (2.5mM; Takara, #R050B), 1μl Partial R1 – CTACACGACGCTCTTCCGATCT (10μM), 1μl T5′ PCR Primer IIA – AAGCAGTGGTATCAACGCAGAG (10μM), cDNA (5-10ng), nuclease-free water (33μl-volume cDNA), 1μl PrimeSTAR GXL DNA Polymerase (Takara, #R050B) was made. The thermocycler was programmed as follows: 98°C for 30sec, 9cyles of 98°C for 10sec, 65°C for 15sec and 68°C for 8min, 1cycle of 68°C for 10min. The concentration of the resulting product was assessed, and if it fell below 10 ng/μl, additional cycles were carried out as needed. The pooled amplified cDNA was cleaned up with 0.6x AMPure XP or SPRIselect beads and taken up in 50μl 5x diluted Elution Buffer (EB; Qiagen) in nuclease-free water (Thermo Fisher).

We optimised scRaCH-seq so that it can also be applied to 10x Genomics multiome cDNA. A mixture containing 2μl cDNA (2ng), 25μl 2x KAPA HiFi master mix (Roche, #KK2602), 1μl Partial R1 – CTACACGACGCTCTTCCGATCT (10μM), 1μl TSOp1 – TGGTATCAACGCAGAGTACATGGG (10μM), 21μl EB buffer was made. The thermocycler needs to be programmed as follows: 98°C for 3min, 8 cycles of 98°C for 15sec, 64°C for 20sec and 72°C for 7min, and 1 cycle of 72°C for 10min.

#### Hybridization

The amplified cDNA sample (1.5-2μg) was dried using a DNA vacuum concentrator (speed vac) together with 1μl of 1000μM IDT Blocking Oligos: Poly A: TTTTTTTTTTTTTTTTTTTTTTTTTTTTTT/3InvdT/, PR1: CTACACGACGCTCTTCCGATCT, SO: AAGCAGTGGTATCAACGCAGAGTAC. Subsequently, the dried sample was taken up in the following mixture: 8.5μl of 2x Hybridization Buffer, 2.7μl Hybridization Buffer Enhancer (xGen IDT, cat#1080577) and 1.8μl nuclease-free water. The sample was incubated at 95°C for 10min to denature the cDNA. To this denatured sample, 4μl of custom probe panel was added and incubated at 65°C for 16 hours to facilitate probe hybridization with the cDNA.

#### Probe Pulldown

We used the xGen IDT Lockdown Hybridization and Wash Kit. The sample was added directly to the dried washed beads (100μl of Dynabeads M-270 Streptavidin) and incubated at 65°C for 45min. Every 10min the sample and beads were mixed. The captured cDNA was then thoroughly washed, and the bead plus sample mixture was resuspended in 50μl EB.

#### Amplification of Captured cDNA Sample

The captured cDNA with beads was amplified using LA Taq DNA Polymerase Hot-Start (Takara, # RR042B) in 4x50μl reactions. For a 50μl reaction, a mixture containing 5μl 10x PrimeSTAR buffer, 4μl dNTP (2.5mM), 1μl FPSfilA: ACTAAAGGCCATTACGGCCTACACGACGCTCT TCCGATCT (10μM), 1μl RPSfilBr: TTACAGGCCGTAATGGCCAAGCAGTGGTATCAACGCAGAGTA (10μM), 12.5μl cDNA on beads, 26.1μl nuclease-free water, 0.4μl LA Taq DNA Polymerase was made. The thermocycler was programmed as follows: 95°C for 2min, 12cyles of 95°C for 20sec, and 68°C for 10min, 1cycle of 72°C for 10min. After amplification, the cDNA was cleaned up with 0.6x AMPure XP or SPRIselect beads and taken up in 30μl EB.

### Nanopore library preparation and sequencing

The Oxford Nanopore Technology (ONT) SQK-PCB111.24 kit was used to index the samples. Since the cDNA will have the ONT overhang, we started from the “Selecting for full-length transcripts by PCR” step in the protocol with 5μl (1ng/μl) scRaCH-seq library, 0.75μl Unique Barcode Primer, 6.75μl Nuclease-free water and 12.5μl 2x LongAmp Hot Start Taq Master Mix. The thermocycler was programmed as follows: 95°C for 30sec, 5 cycles of 95°C for 15sec, 62°C for 15sec and 65°C for 1min, and 1 cycle of 65°C for 6min. After the PCR, the sample was cleaned up with 0.6x beads. For the BTK and SF3B1 gene capture, 21 samples were pooled together and sequenced on PromethION R9.4.1 flow cell (**Figure 1**).

The scRaCH-seq libraries can also be sequenced on a PromethION R10.4.1 flow cell and using the SQK-NBD114.24 Native Barcoding Kit 24 to index the samples and pool them together. 200 fmol of the scRaCH-seq library was used as input.

### Short-read data analysis

#### Data processing

We used *cellranger* (v5.0.0) and *bcl2fastq* (v2.19.1) to pre-process our short-read sequencing data. The percentage of mitochondrial gene counts was calculated using the ‘PercentageFeatureSet’ function in *Seurat* (v4.0.5)(18). This calculation targeted genes starting with the regex pattern “MT-”. For each sample, the ‘isOutlier’ function from *Scater* (v1.20.0)(19) was used to identify low-quality cells. These were defined as cells deviating by more than 3 median absolute deviations (MADs) from the median in terms of unique molecular identifiers (UMIs, both higher and lower), detected gene numbers (higher and lower), and mitochondrial gene expression (higher). Cells identified as low-quality were then excluded from further analysis. Library size normalization and log(x + 1) transformation were performed using the ‘NormalizeData’ function in *Seurat*. This step was followed by the identification of highly variable genes (HVGs) using the ‘FindVariableFeatures’ function in *Seurat*, adhering to default parameters. These HVGs were then scaled and centred with the ‘ScaleData’ function in *Seurat*.

#### UMAP by gene expression

We performed principal component analysis (PCA) by applying the ‘RunPCA’ function in *Seurat* to scaled HVGs (n=2000) with the default parameter settings. To remove the unwanted variability due to batch effects, we applied *Harmony* (v0.1.0) for batch correction(20). For cell clustering analysis, we employed the shared nearest neighbour (SNN) method implemented in *Seurat*. To construct the SNN graph, we used the ‘FindNeighbors’ function with the first 20 Harmony corrected PCA embeddings and k.param = 20. For visualization, we generated a Uniform Manifold Approximation and Projection (UMAP) using the ‘RunUMAP’ function with the first 20 *Harmony*-corrected PCs. To identify cell clusters, we executed the ‘FindClusters’ function on the SNN graph, employing the original Louvain algorithm with default parameters. To identify the doublets, we removed the cluster that exhibited the expression of multiple immune cell lineage markers and high RNA contents. Finally, we applied the same *Seurat* functions on the filtered data to re-cluster cells and update the UMAP.

#### Single-cell differential expression analysis

To perform single-cell differential expression analysis, we employed the ‘FindMarkers’ function of *Seurat*, using the parameters configured as test.use = “MAST”, logfc.threshold = 0 and min.pct = 0.

#### Pseudo-bulk differential expression analysis

The count matrix for gene expression was aggregated using the ‘aggregateAcross’ function incorporated in *Scater*. Subsequently, the Likelihood ratio test, implemented in *edgeR* (v3.34.0) was applied to identify differentially expressed genes (DEGs)(21).

#### Gene set enrichment analysis

We downloaded the C2 canonical pathways and Hallmark pathway collections using *msigdbr* (v7.5.1)(22). All genes were then ordered in a decreasing fashion based on their log-fold changes. This ordered list of genes was then used as input for the ‘GSEA’ function in *ClusterProfiler* (v4.4.4), which was tested against the gene set collections using default parameters(23).

### scRaCH-seq data analysis

We developed a customized *FLAMES* (v1.3.4) pipeline to conduct the scRaCH-seq analysis (**Figure 1**). A detailed description of the original *FLAMES* pipeline can be found in the Methods paper that introduces single-cell Full-Length Transcriptome sequencing (scFLT-seq)(17).

#### Data preprocessing (STEP 01)

We base called the raw data of nanopore sequencing using the *Guppy* software (v3.1.5) with the “dna_r9.4.1_450bps_sup_prom.cfg” configuration, resulting in the generation of fastq files. Subsequently, we used the “find_barcode’’ function in *FLAMES* to demultiplex the fastq files, which was accomplished by cross-referencing the cell barcodes identified in the corresponding scRNA-seq data. We allowed 2 base pairs of edit distance for the cell barcode matching.

#### Read integrity check (STEP 02)

We added an extra step of reads integrity check to the *FLAMES* pipeline. The demultiplexed reads underwent an examination to confirm the presence of all essential components. These components include a cell barcode, a unique molecular identifier (UMI), a template switch oligo (TSO) sequence, and a poly-A tail at the end of the transcript.

#### Point mutation calling (STEP 03)

The reads were first aligned to the human genome GRCh38 (downloaded from Gencode) using the *FLAMES* function “minimap2_align”. We then used the mutation calling function in *FLAMES* (v0.1) with the default parameters to identify the point mutations, by comparing the pile-up reads against the gene of interest. For SF3B, the gene region Chr2: 197,388,515 – 197,435,079 from the human genome GRCh38 was selected. For each sample, the positions with alteration frequencies ranging from 10% to 90% were identified as mutations. The allele frequency of identified mutations was then counted and subsequently summarized to generate a mutation count matrix, with the rows corresponding to mutations and the columns named after the cell barcodes.

#### Deletion calling (STEP 03)

We incorporated an additional step specifically to detect multiple base pair deletions in genes of interest. We first aligned the reads which passed the integrity check to the human genome GRCh38, using *minimap2* with parameters set as “-ax splice --junc-bonus 1 -k14 --secondary=no --junc-bed”(24). We then compared the pile-up from the reads to the reference by checking the alignments at the specific chromosomal position. For the SF3B1 6bp deletion previously identified by WES, read alignments were checked at the Chr2:197,402,104 - 197,402,109 position to identify the presence of the 6bp deletion. The reads with the multiple bp deletion were merged by UMIs and subsequently counted for each cell to generate a count matrix, with rows representing the SF3B1 6bp-deletion/WT and column names corresponding to the cell barcodes.

#### Isoform detection (STEP 04)

We employed the *FLAMES* function “find_isoform” to summarize the alignment for making isoform assembly. The isoforms that exhibited <10bp variance in their splicing junctions and <100bp variance in their start or end sites were merged. Following this, we re-aligned the reads that passed the integrity check to the isoform assembly to generate the isoform count matrix, with the rows corresponding to isoforms and the columns named after the cell barcodes.

#### Data integration (STEP 05)

We combined the isoform and mutation count matrices with the processed single-cell short-read data to construct a comprehensive multi-assay *Seurat* object, using the ‘CreateAssayObject’ function. The scRaCH-seq data were used to identify cell groups with specific isoforms or mutations, followed by short-read differential expression analysis between these groups.

#### Differential transcript usage analysis

For scRaCH-seq data, we merged the per-cell counts of target gene transcripts by samples to make pseudo-bulk transcript data. We calculated the frequency of different transcripts for our genes of interest SF3B1 and BTK respectively and then displayed the top 5 transcripts with the highest frequencies across all samples for both target genes. For scFLT-seq data, cells from patients harbouring SF3B1 mutations/deletions were combined per sample to make pseudo-bulk transcript data for the SF3B1-mutated group. Cells of wild-type screening/venetoclax-relapsed/healthy donor samples were merged to construct screening-wt/relapsed-wt/healthy-wt pseudo-bulk transcript data for the SF3B1-wild type group. We then applied the function “diffSpliceDGE” in *edgeR* to identify the differential transcript usage between groups (SF3B1-mutated, SF3B1-wt) using pseudo-bulk data as replicates.

## Results

### Assessing data quality and probe-capture efficiency

After scRaCH-seq was performed, a total of 43,762,097 raw reads were acquired through Nanopore sequencing across 21 samples, with a read length distribution spanning from 0 to 4,000bp (**Figure 2A**). The *FLAMES* demultiplexing process was subsequently able to recover 28,573,271 reads that could be attributed to specific cells. Despite an overall 35% read loss during this step, the length distribution of the retained reads remained unaltered (**Figure 2A**). After demultiplexing, sufficient reads (∼1,000,000) were preserved for each sample (**Figure 2B**). We also tested Flexiplex(25) and the reads retained for downstream processing were similar (data not shown). Next demultiplexed reads without TSO sequences and poly-A tails were discarded following the quality control step which eliminated around 49% of reads, primarily those with lengths < 1500bp (**Figure 2A**). A total of 14,442,490 reads were retained for subsequent read alignment and feature counting. Moreover, scRaCH-seq and our *FLAMES* pipeline effectively captured the majority of cell barcodes identified by *CellRanger* in scRNA-seq data (**Figure 2B**). 90% of the reads retained aligned successfully to the targeted genes SF3B1 and BTK (**Figure 2B**). Duplicated reads were collapsed using UMIs. In concordance with on-target reads, a high number of UMIs were associated with SF3B1 and BTK, with the majority of captured cells harbouring the target genes (**Figure 2C**). The captured off-target genes include HSPD1, which is near SF3B1, and TIMM8A, located in close proximity to the BTK gene (**Figure 2D**). These were some read-through transcripts but also background reads. Per-cell UMI counts for the target genes in the corresponding short-read data were evaluated, revealing a higher per-cell UMI capture in the scRaCH-seq data for SF3B1 and BTK (**Figure 2E**). In two separate scRaCH-seq experiments targeting 17 BCL2 family genes in CLL cells and 24 genes commonly mutated in acute myeloid leukaemia (AML) cells, similar UMI counts were observed compared to short-read data **(Supplementary Figure 2)**. The AML sample was processed using the 10x Genomics Chromium single-cell 3’ kit. These findings indicate the high efficiency of scRaCH-seq in capturing the transcripts of target genes observed in matched single-cell 5’ and 3’ short-read datasets via long-read sequencing.

**Figure 2.**
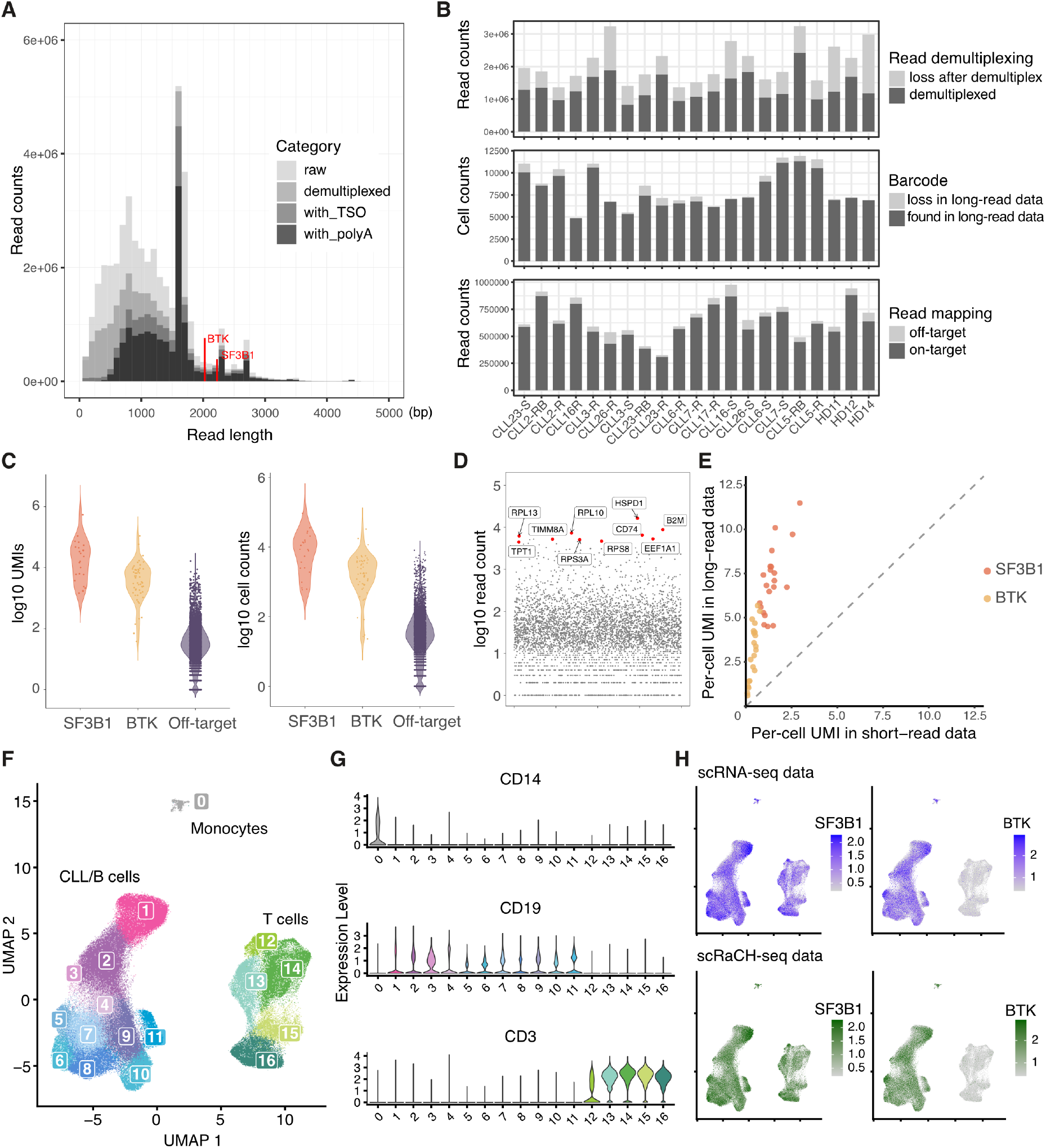
scRaCH-seq is efficient in capturing enriched genes of interest. **A)** A graph showing the losses in read counts due to demultiplexing and integrity checks, depicting the distribution of read lengths ranging from 0 to 5,000 base pairs. The canonical isoform of BTK (2,027 bp) and SF3B1 (2,225 bp) are marked on the plot. The four shades of grey, ranging from light to dark, correspond to raw reads, demultiplexed reads, reads with TSO, and reads with also a poly-A end. **B)** Bar plots showing the loss of reads after demultiplexing (top panel), barcodes detected in long-read sequencing data (middle panel) and counts of off-target and on-target reads for each sample (bottom panel). **C)** Violin plot showing the log UMI counts of BTK (red), SF3B1 (orange) and off-target genes (blue) detected by scRaCH-seq (left panel). Violin plot showing the log counts of cells possessing UMIs of BTK (red), SF3B1 (orange) and off-target genes (blue) (right panel). **D)** Dot plot showing the log counts of off-target genes with the top 10 off-target transcripts specifically marked. **E)** Dot plot showing the per-cell UMIs of SF3B1 (red) and BTK (orange), detected by scRNA-seq (X-axis) and scRaCH-seq (Y-axis) for all samples. **F)** UMAP projection of peripheral blood mononuclear cells from CLL patients or healthy donors and clustering based on short-read gene expression. **G)** Violin plot showing the expression of CD14 (monocyte marker), CD19 (B/CLL cell marker) and CD3 (T cell marker) per cell cluster (Figure 2F). **H)** UMAP projection of SF3B1 (left) and BTK (right) gene-level expression detected by scRNA-seq (purple) or scRaCH-seq (green).

Next, we compared the gene expression of *SF3B1* and *BTK* in the scRaCH-seq data with the short-read dataset. Similar to the short-read gene expression data, scRaCH-seq showed that *BTK* expression was detected in the CLL/B cell population, whereas *SF3B1* was expressed across all cell types. (**Figure 2F-H**)

### Isoform detection by scRaCH-seq

Given that scRaCH-seq operates by capturing genes with probes, it enables the exploration of isoform usage for the targeted genes. In our dataset, we incorporated the CLL and B cells from matched samples from CLL patients at diagnosis, after venetoclax relapse, in patients who relapsed and subsequently received a BTK inhibitor (BTKi) and healthy donor samples(16). The primary BTK isoform (BTK-201, ENST00000308731) captured by scRaCH-seq is a protein-coding isoform with 19 exons, exhibiting high expression across all samples (**Figure 3A**). The other four transcripts among the top 5 captured isoforms by scRaCH-seq are novel isoforms not catalogued previously (**Figure 3A**). Transcripts #2 and #3 lack the last four exons compared to BTK-201, with transcript #2 having a 16-base pair longer exon 10 compared to transcript #3. Novel transcript #4 lacks exon 10 compared to the BTK-201 transcripts. While all five BTK transcripts detected by scRaCH-seq have a poly-A tail in the 3’ UTR, transcript #5, comprising only three exons, has a different starting point at the 5’ UTR. Consequently, it remains uncertain whether BTK transcript #5 is an artefact resulting from truncation at the 5’ end. No evidence of differential BTK transcript usage was found across the different sample groups (**Figure 3A**).

**Figure 3.**
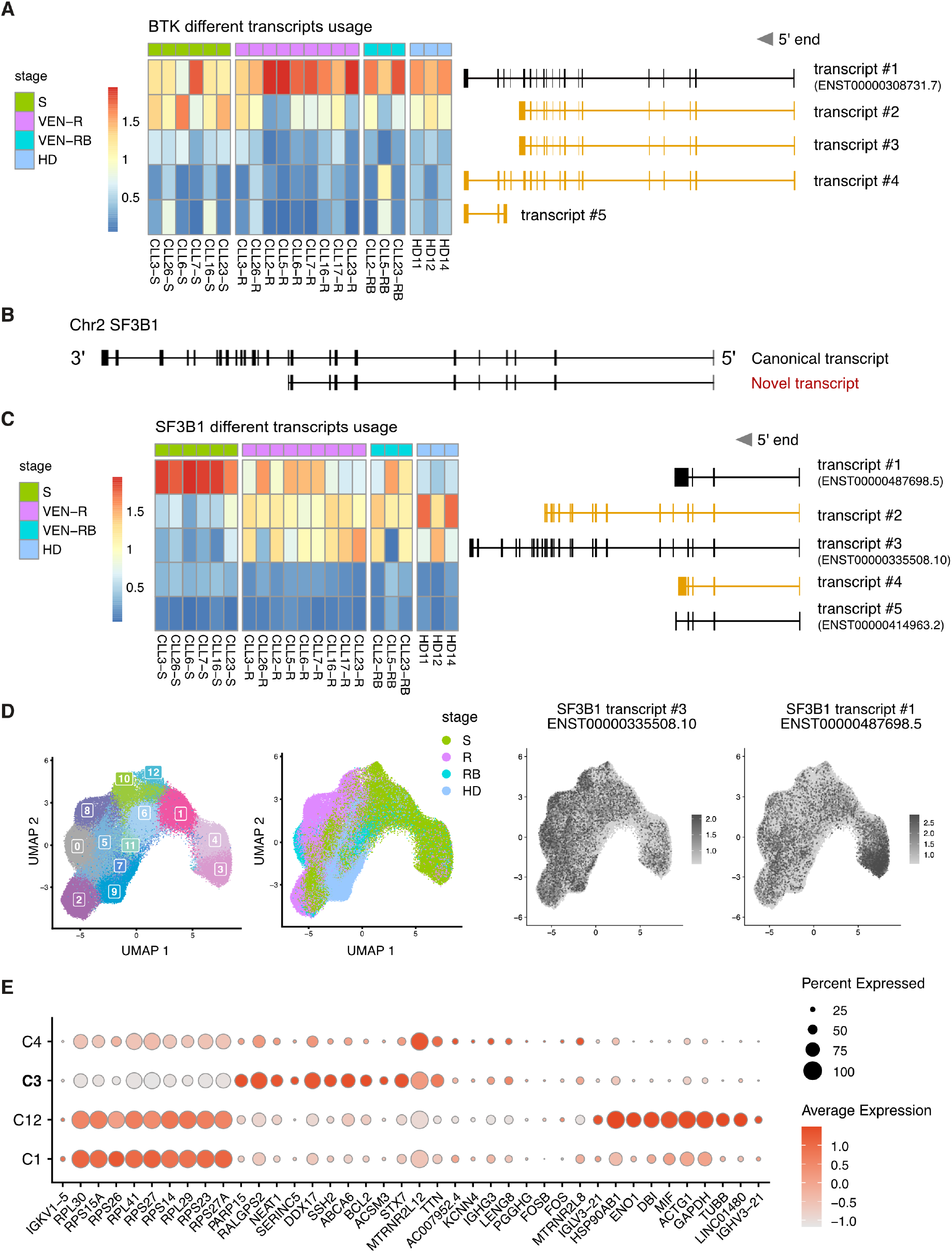
scRaCH-seq reveals different SF3B1 isoform usage by CLL cells from patients undergoing different treatments. **A)** Heatmap showing BTK isoform usage (rows) per sample (columns) in the B/CLL cells. The top 5 BTK transcripts are illustrated on the right with novel transcripts highlighted in yellow. Samples are grouped in screening (S; green), venetoclax relapsed (VEN-R/R; pink), venetoclax relapsed and subsequently on BTKi (VEN-RB/RB; blue) and healthy donor (HD; purple). **B)** Illustration of the dominant SF3B1 transcript identified and characterized by the absence of the last 14 exons in the 3’ end (highlighted in red). **C)** Heatmap showing SF3B1 isoform usage (rows) per sample (columns) in the B/CLL cells. The top 5 SF3B1 transcripts are illustrated on the right with novel transcripts highlighted in yellow. **D)** UMAP projection of CLL/B cells from CLL patients or healthy donors and clustering based on short-read gene expression. The samples from CLL patients at screening (S; green), venetoclax relapsed (R; pink), venetoclax relapsed and subsequently on BTKi (RB; blue) and healthy donors (HD; purple) are highlighted in the UMAP. The expression of the SF3B1 transcript #1 and #3 are overlaid as dark grey dots in the UMAP (Figure 3B). **E)** Dot plot showing marker genes that distinguish the CLL cells present in the 4 clusters shared by the screening samples (C1, C3, C4 and C12 in Figure 3C). Cluster 3 with SF3B1 transcript #1 usage is indicated in bold.

In our investigation of the SF3B1 isoforms, a previously unidentified SF3B1 transcript emerged as the predominant form, characterized by the absence of the last 14 exons in the 3’ end (**Figure 3B)**. Given that this transcript is not the canonical SF3B1 transcript expressed in the majority of cells, including healthy donor samples, we undertook a comprehensive assessment to determine its authenticity. This transcript does not appear to be an artefact induced by Nanopore sequencing, since it initiates at exon 1 and concludes with a poly-A tail. To further validate its presence, we examined the distribution of SF3B1 reads in scRaCH-seq data and compared it to matched bulk RNA-seq and scFLT-seq data for sample CLL2-RB. A consistent pattern in the SF3B1 read distribution between scRaCH-seq and scFLT-seq was observed, with a higher coverage of exons 1–11 compared to that of exons 12–25 (**Supplementary Figure 3A**). However, when comparing single-cell sequencing data (scRaCH-seq and scFLT-seq) to bulk RNA-seq, bulk RNA-seq data exhibited a relatively stable read coverage across all exons (**Supplementary Figure 3A)**. This hints at a potential bias introduced by the 10x Genomics protocol into the composition of SF3B1 transcripts. Upon closer examination, we discovered that 86% of single-cell reads mapped to exon 11 contained three cytosine (C) to thymine (T) mismatches following the drop in read coverage, indicating that these reads were created due to the unexpected binding of 10x poly-A primers (**Supplementary Figure 3B)**. These artefacts were subsequently removed from the scRaCH-seq data, resulting in the elimination of the novel SF3B1 transcript which lacks the last 14 exons (**Supplementary Figure 3C**).

After removal of the artefacts, high expression levels of SF3B1 transcript #1 (SF3B1-211, ENST00000487698), transcript #2 (a novel transcript containing exons 1 – 16) and transcript #3 (SF3B1-201, ENST00000335508) were consistently observed across multiple samples (**Figure 3C**). The expression of transcripts #4 (a novel transcript) and #5 (SF3B1-204, ENST00000414963) was relatively low (**Figure 3C**). In contrast to BTK, a clear difference in SF3B1 isoform usage was evident between CLL cells from patient samples at screening and those from other CLL stages or healthy donor B cells (**Figure 3C**). Specifically, SF3B1 transcript #1 was highly expressed in screening samples, whereas transcripts #2 and #3 were highly expressed in venetoclax-relapsed and healthy donor samples. In contrast to the CLL and B cells, SF3B1 transcript #3 was the dominant transcript expressed by the T cells from the CLL patients at different stages and healthy donors (**Supplementary Figure 4**). To investigate why the screening samples exhibit higher expression of transcript #1, we linked isoform usage with gene expression. The CLL and B cells from all samples were isolated and re-clustered, revealing distinct clusters of the different stages, including one B-cell cluster (C9), four screening clusters (C1, C3, C4, C12), three relapsed clusters (C0, C2, C8), and five clusters shared by screening and relapsed CLL cells (C5, C6, C7, C10, C11) (**Figure 3D**). Interestingly, SF3B1 transcript #1 was specific to the screening CLL cells from cluster 3 (**Figure 3D**). Meanwhile, transcript #3, the canonical SF3B1 isoform, was expressed by all CLL and B cells (**Figure 3D**). Subsequently, single-cell differential gene expression analysis between all the screening clusters (C1, C3, C4, C12) was performed. For this analysis, other clusters were excluded to eliminate confounding factors related to treatment stages. CLL cells in C3 exhibited a high expression level of DDX17, a gene involved in almost all RNA metabolism processes(26) and PARP15, a negative regulator of transcription (27) (**Figure 3E**).

### Calling SF3B1 mutations using scRaCH-seq data

Besides isoform detection, scRaCH-seq has the ability of linking the mutation status to cell state and differential gene expression. For the identification of SF3B1 mutations, we aggregated the scRaCH-seq data from all the samples, revealing many SF3B1 transcripts with single-nucleotide polymorphism (SNP) (**Figure 4A**). The three most prominent SNPs were #5, #6, and #8. While SNP #6 and #8 exhibited a low frequency (∼20%) across all samples, SNP #5 was unique to four specific samples. SNP #5 corresponds to the SF3B1 K700E mutation (Chr2:197402110 T→C), a mutation frequently observed at a sub-clonal level in CLL patients(28-30). The K700E mutation can interrupt the recruitment of SF3B1 to the correct 3’ end branch site and consequently result in alternative splicing (31). We identified the SF3B1 K700E mutation in CLL3 and CLL26 in both screening (S) and venetoclax-relapsed (R) samples (**Supplementary Figure 5A**). This mutation was also confirmed by the WES data from these samples(16). In scRaCH-seq, the K700E mutation was consistently observed with an overall frequency of 45.0% across the four samples, encompassing 87,674 UMI counts at the location of the SF3B1 K700E mutations and 48,224 UMI counts containing the reference base (Thymine) (**Figure 4B**). Although SNP #1 and #3 were specific to four samples and occurred at a high frequency, the overall UMI coverage for these altered transcripts was low (**Figure 4A-B**). A comparison of scRaCH-seq efficiency in capturing the SF3B1 K700E mutation with whole transcriptome short-read sequencing (scRNA-seq) and long-read sequencing (scFLT-seq) revealed that scRaCH-seq significantly increased the capture of cells carrying the mutation and enriched for transcripts (**Figure 4C**).

**Figure 4.**
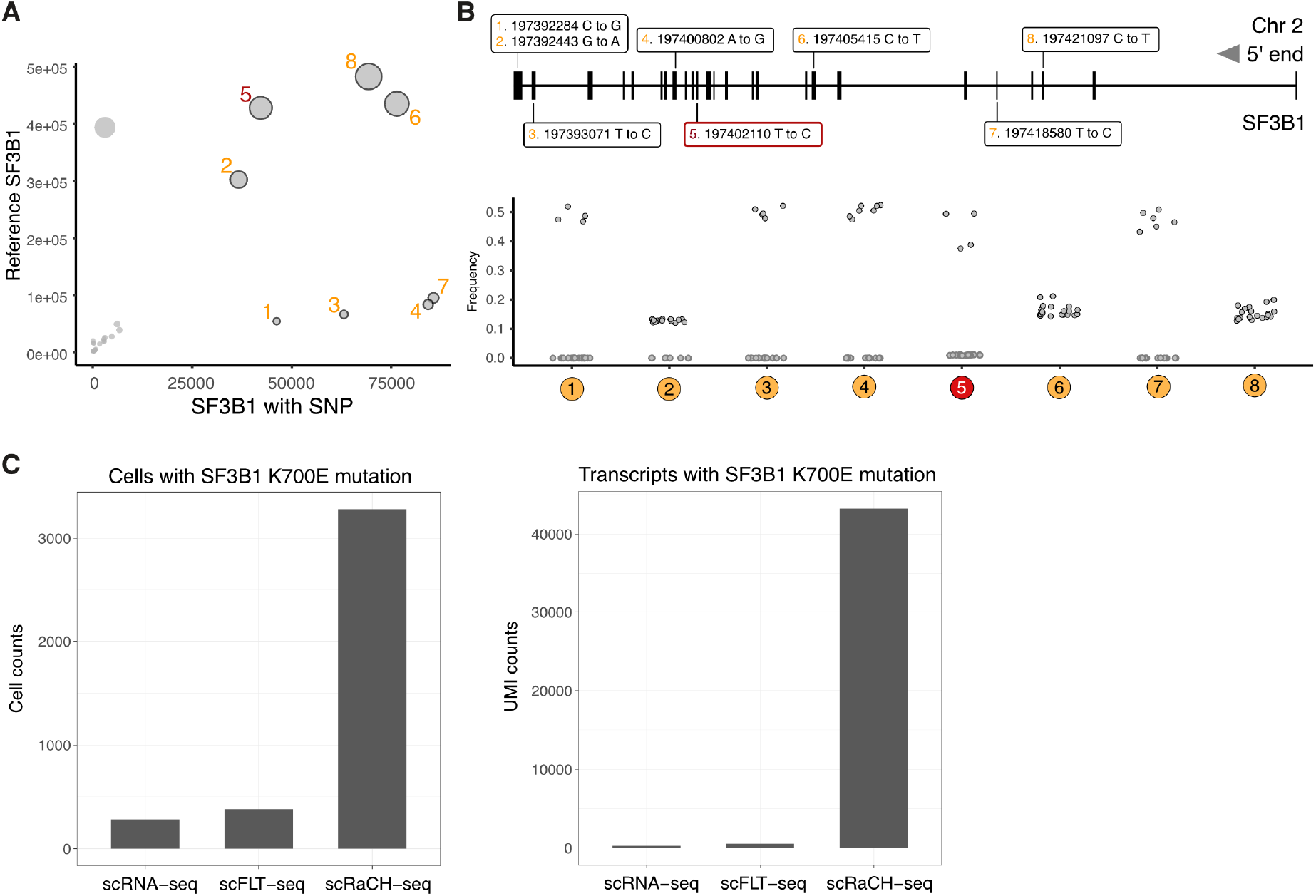
Mutation calling with scRaCH-seq and FLAMES. **A)** Dot plot showing the SF3B1 point mutations identified using FLAMES (left panel). After consolidating reads with UMIs, the reads from all the samples were aligned to the GRCh38 reference genome. The size of each dot in the plot corresponds to the proportion of altered transcripts. **B)** A graphical representation highlights the precise locations of these high-frequency (>10%) alterations within the SF3B1 gene. The bottom graph shows the frequency of altered SF3B1 transcripts in each sample. **C)** Bar plots showing the cells with a SF3B1 K700E mutation (left panel) or transcripts (UMIs) with SF3B1 K700E mutation (right panel) detected by short-read scRNA-seq, long-read scFLT-seq or long-read scRaCH-seq.

Subsequently, we isolated cells harbouring SF3B1 K700E mutant reads from the scRaCH-seq data and visualized their distribution on the UMAP layout, which contained all patient sample cells (**Figure 5A**). Most cells with the mutant reads clustered within the CLL-cell group, consistent with the findings from WES. However, a small number of cells with mutant reads were detected within the non-CLL clusters, including the B-cell cluster from a healthy donor, T-cell cluster, and monocyte cluster (**Figure 5A and Supplementary Figure 5B**). Given that WES established the absence of SF3B1 mutations in non-CLL cells, we investigated whether these non-CLL cells with SF3B1 mutant reads might be attributed to false positives induced by doublets. Comparative analysis of UMI counts and feature counts revealed that non-CLL cells with SF3B1 mutant reads exhibited similar values to CLL cells with SF3B1 mutants (**Figure 5B**). Moreover, expression of multiple lineage markers within the non-CLL cells with SF3B1 mutant reads was not observed, suggesting that these cells were not doublets (**Figure 5B**). In the SF3B1 wild-type samples confirmed by WES, we identified altered reads with a read fraction of less than 1% (**Figure 5C and Supplementary Figure 5B)**, indicative of reads containing sequencing errors(32). To mitigate these false positives in both wild-type samples and non-CLL clusters, we established a threshold based on the abundance of SF3B1 mutant transcripts detected per cell. Setting the threshold at more than 1 SF3B1 mutant transcripts with different UMIs, resulted in the exclusion of 80.7% of cells with SF3B1 mutant transcripts in wild-type samples (**Figure 5C**). Furthermore, by raising the threshold to more than 2 SF3B1 mutant transcripts, 93.4% of false positives in wild-type samples were successfully eliminated (**Figure 5C**). Consistent with false positives observed in wild-type samples, most SF3B1-mutant cells in non-CLL clusters exhibited low counts of altered reads (**Figure 5D**). Applying the threshold of more than 2 mutant reads per cell for the identification of a true positive mutant SF3B1 cell resulted in the removal of most false positives in non-CLL clusters (**Figure 5E**). Employing the threshold of more than 2 mutant transcripts based on abundance was then established as a standard quality control step to eliminate false positives during mutation calling. The transcript with altered SF3B1 #1 was observed in matched samples from CLL3 and CLL5 (**Supplementary Figure 5C**). This alteration was observed in both CLL clusters and non-CLL clusters in the UMAP (**Supplementary Figure 5D**). These observations suggest that the C-to-G alteration in the SF3B1 3’ UTR is likely a SNP present at a baseline level in specific patient samples. Notable, even with the threshold in place, we identified SNP #8, a C→T alteration at Chr2:197421097, in all samples and across all cell types (**Supplementary Figure 5E-F**).

**Figure 5.**
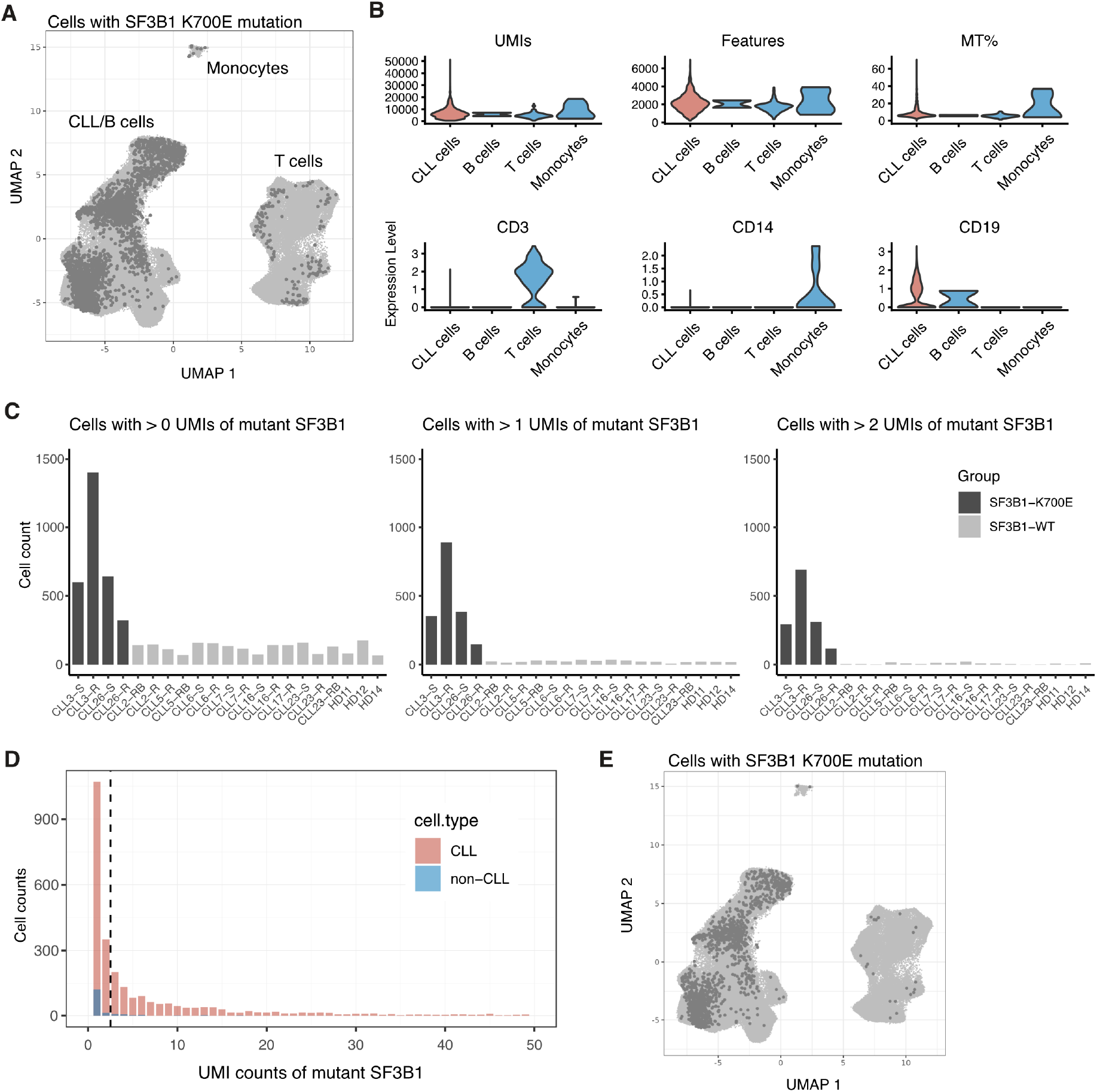
The SF3B1 K700E mutation is detected by scRaCH-seq. **A)** UMAP projection of cells carrying the SF3B1 K700E mutation (dark grey). **B)** Violin plot showing the UMI counts, gene counts (Features), expression of mitochondria genes (MT%), expression of CD3 (T cell marker), CD14 (monocyte marker) and CD19 (B cell marker) per cell type cluster. **C)** Bar plot showing the abundance of mutant transcripts. Left plot showing cells per sample with >0 transcripts (UMI) of mutant SF3B1 K700E. Middle plot showing cells per sample with >1 transcripts (UMIs) of SF3B1 mutation. Right plot showing cells per sample with >2 transcripts (UMIs) of SF3B1 mutation. Samples with confirmed SF3B1 K700E mutation by WES are highlighted in dark grey. **D)** Bar plot showing the distribution of the abundance of SF3B1 K700E mutation detected in CLL (pink) and non-CLL (blue) cells. The vertical dashed line indicates the criterion of >2 SF3B1 mutation transcripts. **E)** UMAP projection of cells carrying >2 SF3B1 K700E mutation transcripts (dark grey).

### Detecting the 6-base pair deletion in SF3B1 among CLL cells

In addition to the SF3B1 K700E point mutation, WES also confirmed the presence of a 6-bp deletion (6bp-del) at Chr2:197402104:197402109, resulting in an SF3B1 KVR700**R mutation in CLL17 at a subclonal level(16). The original *FLAME*S pipeline was only designed to identify point mutations, so to address this limitation, we developed a supplementary script dedicated to detect the specific deletion directly in the fastq files of all samples. To minimize potential false positives, we applied the same criterion as we did for point mutation calling, necessitating at least two transcripts with the deletion per cell across all samples. Notably, cells with the 6bp deletion were exclusively observed in sample CLL17-R (**Figure 6A**), with a predominant clustering within the CLL group (**Figure 6B**), aligning with WES findings and underscoring the reliability of the deletion counting method. Subsequently, single-cell differential gene expression analysis was performed to identify gene expression changes between CLL cells with the SF3B1 deletion and wild-type cells across all venetoclax-relapsed (R) samples. The analysis revealed a total of 377 DEGs with a |logFC| > 0.5 and adjusted *p*-value < 0.05. Among these, 152 genes showed significant up-regulation, while 225 genes were down-regulated in CLL cells with the KVR700**R mutation (**Figure 6C**). Similarly, we identified 147 differentially expressed genes (FDR < 0.05) between K700E mutant and wild-type CLL cells, comprising 28 up-regulated genes and 119 down-regulated genes in SF3B1 K700E CLL cells (**Supplementary Figure 6A**). Given the sub-clonal nature of the SF3B1 KVR700**R mutation in CLL17, DE analysis between mutant and wild-type SF3B1 CLL cells of CLL17 was also conducted. Surprisingly, we did not observe any DEGs (**Figure 6D**) This implies that the identified DEGs between cells carrying the SF3B1 6-bp deletion and wild-type cells are likely relapsed patient-specific rather than mutation-driven.

**Figure 6.**
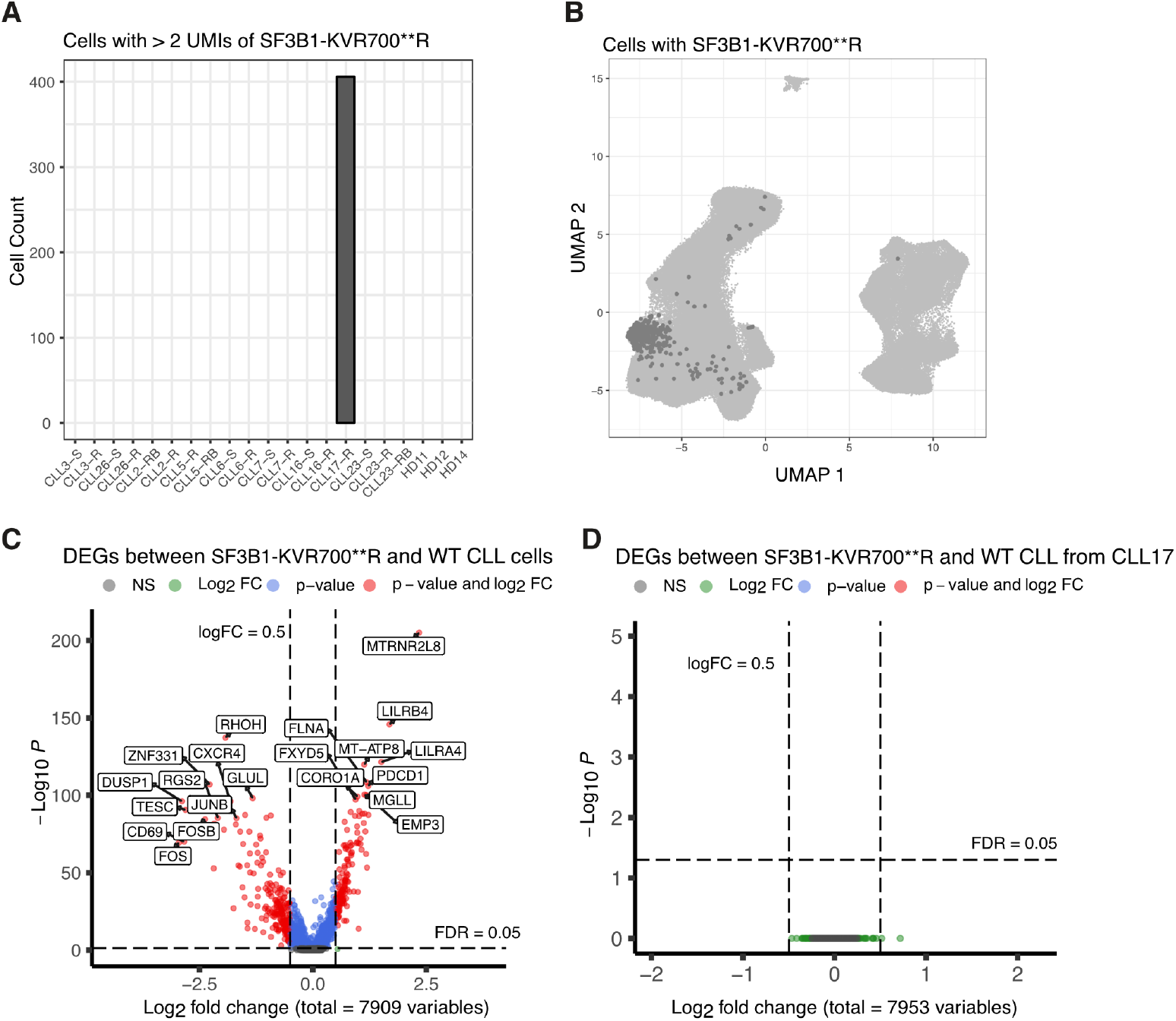
The SF3B1 KVR700**R alteration is detected by scRaCH-seq. **A)** Bar plot showing the cells with SF3B1 6bp-deletion (Chr2:197402104:197402109) across all samples. A criterion of > 2 of SF3B1 with 6bp-del transcripts (UMIs) per cell was employed. **B)** UMAP projection of cells carrying >2 SF3B1 KVR700**R altered transcripts (dark grey). **C)** Volcano plot showing the differentially expressed genes (DEGs; FDR<0.05) between SF3B1 KVR700**R mutant and wild-type CLL cells from all venetoclax relapsed CLL samples. **D)** Volcano plot showing the differentially expressed genes (DEGs; FDR<0.05) between SF3B1 KVR700**R mutant and wild-type CLL cells from CLL17.

### Altered splicing in CLL cells with SF3B1 mutations

It has been reported that the inhibition of RNA splicing can enhance the sensitivity of venetoclax in a mouse model bearing acute myeloid leukaemia (AML)(33). Therefore, the question remains if SF3B1 mutation and downstream altered splicing events can contribute to venetoclax resistance in patients with CLL. The current scRaCH-seq probe panel was designed for two target genes to detect isoform usage and mutation calling. However, scFLT-seq from the same samples offers a broader overview of all genes at the transcript level, albeit with lower read coverage per gene(16). Interrogating the single-cell full-length transcriptomic data showed altered splicing in the CLL cells with SF3B1 mutation including altered ‘3 splicing of the TPT1 gene as an example (**Figure 7A-B**). This is coherent with previous reports(34,35), however, we found no evidence of altered splicing transcripts specific to the relapsed SF3B1 mutated samples (**Figure 7C**). Furthermore, no altered splicing of the BCL2 family genes was observed in SF3B1 mutated CLL cells. This finding, in conjunction with the shared enriched pathways uncovered by scRNA-seq data (**Supplementary Figure 6B-C**), suggests that SF3B1 mutations may not play a significant role in the development of acquired venetoclax resistance in CLL cells.

**Figure 7.**
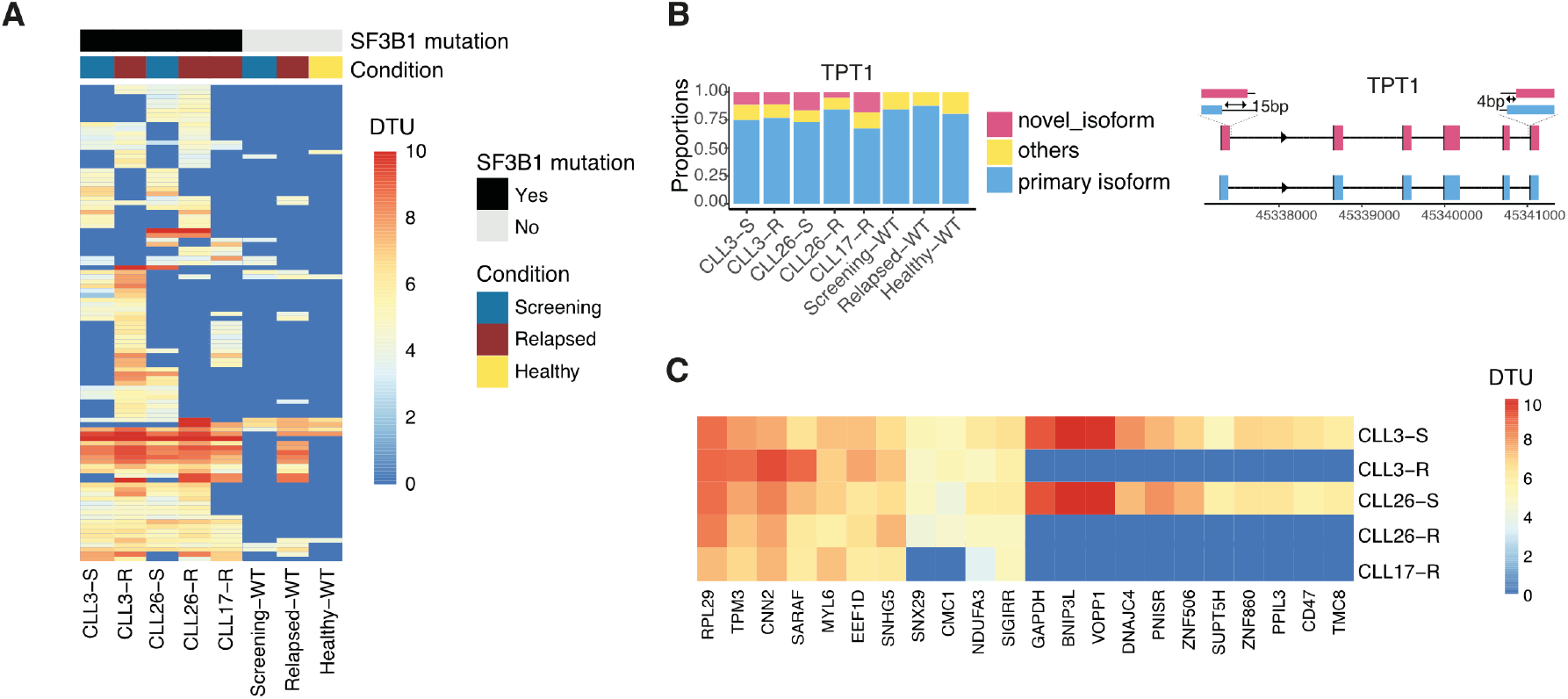
Novel transcript usage in SF3B1 mutated CLL samples. **A)** Heatmap showing differential transcript usage (DTU) compared to primary isoform for genes (rows) per sample (columns). The wild-type SF3B1 CLL cells from patients at screening (blue), at venetoclax-relapsed (red) and the healthy donor B cells (yellow) are combined. **B)** Bar plot showing the novel isoform usage of the TPT1 gene selected from the heatmap in Figure 8A. Novel spliced isoform is highlighted in pink and illustrated in the right panel. **C)** Heatmap showing differential transcript usage (DTU) compared to primary isoform for selected genes (columns) per SF3B1-mutated sample (rows). The genes are selected based on differences between the screening and venetoclax-relapsed groups.

## Discussion

Cell identity and function could be influenced by the alternative splicing of transcripts which will result in a substantial number of transcript isoforms. However, a significant portion of alternative transcripts will not be detected through second-generation sequencing-based single-cell RNA-seq methods due to the short read length and the inherent bias toward 3′ or 5′ ends of the transcripts(11). To address this limitation, we developed scRaCH-seq, demonstrating high specificity and efficiency in capturing targeted long-read transcripts. This method provides an in-depth analysis of transcript usage of genes of interest, adding another layer to the existing single-cell short-read RNA-seq data. Integration of unique barcode primers specific to ONT allowed pooling and efficient sequencing of multiple samples on the Nanopore platform. scRaCH-seq’s capability to capture multiple transcripts simultaneously enhances isoform usage detection and facilitates the discovery of transcripts with mutations, providing a valuable tool across diverse research fields. Our study demonstrated the efficient capture of transcripts <5kb from pre-indexed and stored cDNA, depending on which genes were targeted. While this study focused on the capture of 2 genes that are highly expressed, we demonstrated the efficient capture of the transcripts of 17 and 24 genes, with the potential for larger gene sets without the need for primer optimization. Notably, scRaCH-seq demonstrated concordance with scRNA-seq (10x Genomics) in capturing genes, suggesting its reliability in transcriptomic profiling.

We optimized the *FLAMES* pipeline for scRaCH-seq data by incorporating read integrity checks to eliminate 3’end-truncated reads and setting a quality control threshold of at least two mutant reads to reduce false positives. Additionally, a deletion counting script was created for identifying known deletions. This allowed the detection of the SF3B1 K700E mutation and the 6-base pair deletion in SF3B1 in particular samples, both of which were previously confirmed by WES. Meanwhile, the incidence of false positives stemming from sequencing errors was minimal after the filtering based on the abundance of mutant reads detected in CLL cells. While single-cell DNA sequencing, such as Mission Bio Tapestri, provides insight into clonality upon drug resistance (36), it lacks transcriptome profiling per cell. In contrast, scRaCH-seq offers short-read whole transcriptomic data and mutation status if the mRNA of the targeted gene is expressed. By integrating the short-read scRNA-seq data, we observed multiple significant DEGs between SF3B1 K700E/KVR700**R and wildtype CLL cells including pathways known to be altered in SF3B1 mutated CLL (data not shown)(37). The subclonal nature of SF3B1 KVR700**R in CLL17 allowed us to study the mutation’s impact at a single-cell level. Surprisingly, no DEGs were observed between SF3B1 KVR700**R and wild-type CLL cells from sample CLL17. However, scFLT-seq demonstrated that the SF3B1 KVR700**R mutation increased the expression of novel isoforms in CLL17. Our single-cell data revealed that the altered gene expression between SF3B1-mutated and wild-type CLL cells was primarily driven by patient specificity. This emphasizes the importance of caution when drawing conclusions based on bulk RNA-seq data comparing wild-type and mutant samples from different patients.

Venetoclax is an effective therapy for CLL patients with the potential to induce long remissions. However, we showed that multiple mechanisms resulting in the deregulation of apoptotic genes could occur at a polyclonal level in venetoclax-relapsed patient samples(16). Now by incorporating scRaCH-seq, we can add another layer of information and study the effect of an SF3B1 mutation on venetoclax sensitivity. While it was observed that inhibiting RNA-splicing could enhance venetoclax sensitivity in AML(33), we demonstrated that SF3B1 mutated venetoclax-relapsed CLL cells did not express novel isoforms of the BCL2 family members that could impact venetoclax sensitivity. Interestingly, we observed differential transcript usage of SF3B1 between screening and venetoclax-relapsed samples. SF3B1-211 was found to be specific to a cluster of screening samples but the specific role of this SF3B1 isoform remains unknown. The CLL cells with this SF3B1-211 isoform expressed lower levels of ribosomal genes. These findings suggest a potential connection between the usage of SF3B1-211 transcript and the senescence stage in CLL cells, possibly through its impact on gene transcription activity and protein synthesis.

Lastly, we observed a consistent SF3B1 artefact across all samples in the scRaCH-seq and scFLT-seq data. Comparison of read coverage between scRaCH-seq, scFLT-seq, and bulk RNA-seq data revealed that this artefact resulted from the unexpected binding of the 10x reverse transcriptase primer, rather than being a technical issue with the scRaCH-seq protocol. This unexpected binding of the 10x primers did not impact the results of scRNA-seq since scRNA-seq operates at the gene level, counting all reads aligned to SF3B1. However, it could pose a problem for scFLT-seq and scRaCH-seq, which distinguish reads at the transcript level. Therefore, when identifying novel transcripts in scRaCH-seq or scFLT-seq data, it is advisable to conduct a thorough examination of potential unexpected binding sites of 10x primers within the corresponding gene.

In conclusion, scRaCH-seq provides an innovative strategy for studying long-read transcripts from pre-indexed cDNA, holding promise for advancing gene expression studies and unravelling complex biological processes. Its potential adaptability for PacBio sequencing extends its utility, with the primary advantage lying in linking transcript usage and mutation status at a single-cell level. The approach is both cost-effective, high throughput and flexible, which is achieved by leveraging a wide range of widely used 10x single-cell protocols, making scRaCH-seq applicable for large-scale studies to comprehensively characterize cellular heterogeneity. It holds potential for integration into single-cell spatial data, representing a powerful advancement in single-cell genomics for understanding cellular heterogeneity in development, disease, and other biological processes.

## Data Availability

Single-cell short-read sequencing, whole exome sequencing (WES), and scFLT-seq data are available via the European Genome-phenome Archive (EGA; accession number EGAS00001005815). scRaCH-seq for the *BTK* and *SF3B1* genes are available at EGA50000000158. A data transfer agreement is required, and limitations are in place on the disclosure of germline variants. Any other relevant data are available from the corresponding authors upon reasonable request.

The GitHub link for probe panel design is https://github.com/HongkePn/RaCHseq_Probe_Design. The link for the scRaCH-seq analysis code is https://github.com/HongkePn/scRaCHseq_data_analysis.

## Acknowledgements

We thank Stephen Wilcox, Sarah MacRaild, the WEHI SCORE team and core facilities (Genomics) for their support. Illustrations were created with BioRender.com.

