## Supplementary File for "Single-cell Rapid Capture Hybridization sequencing (scRaCH-seq) to reliably detect isoform usage and coding mutations in targeted genes at a single-cell level"

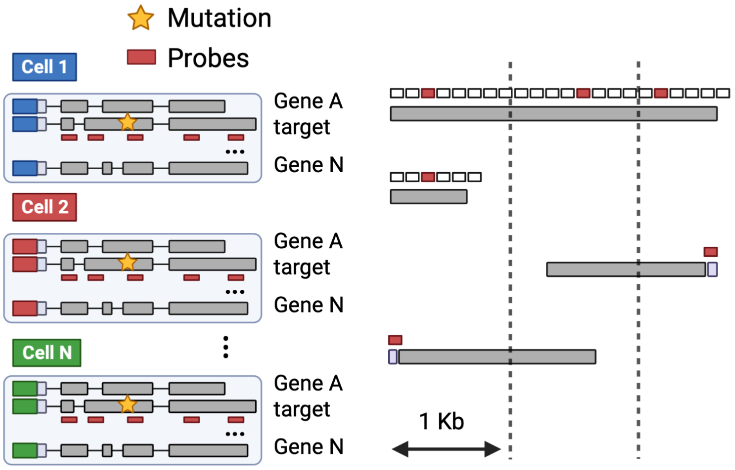


**Supplementary Figure 1. Illustration of the strategy for probe panel design.** Probes were designed to target the gene of interest (left panel). The probes (red and white) were designed to be 120 bp. One probe (red) per kilobase of the target gene exon was selected based on their GC content. If the exons were longer than 1 kilobase, multiple probes were selected per exon. For exons shorter than 120 base pairs, the exons were aligned to create sequences longer than 120 base pairs. This strategy was employed to ensure that each annotated exon was covered by at least one probe.

**Figure 2**

**A**

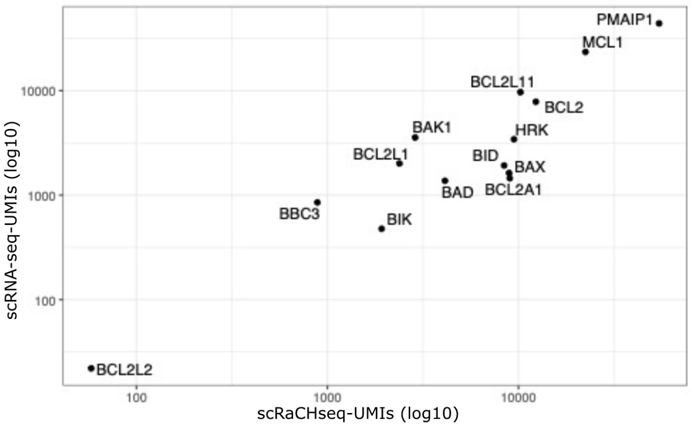


**B**

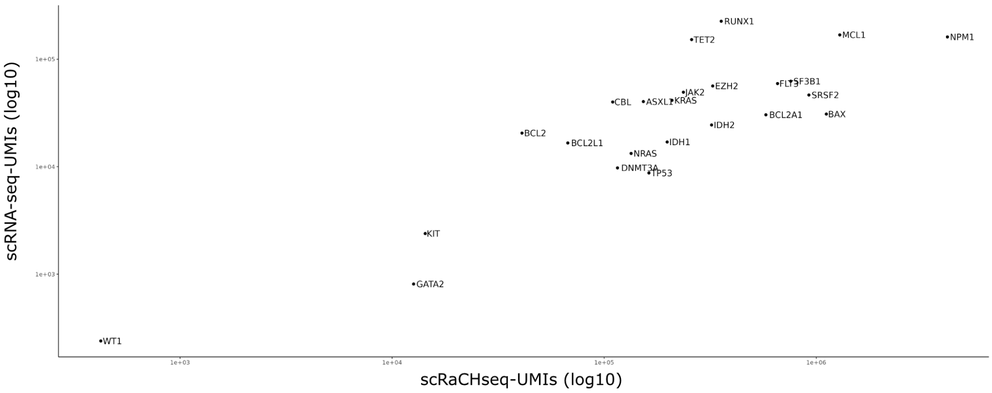


**Supplementary Figure 2. The unique molecule identifiers (UMIs) captured by scRNA-seq and scRaCH-seq are correlated.**

**A)** scRaCH-seq was performed with a probe panel of 292 probes targeting 17 BCL2 family genes. The dots represent the UMIs of target genes detected by scRaCH-seq (X-axis) and scRNA-seq (Y-axis) for a CLL sample.

**B)** scRaCH-seq was performed with a probe panel of 1157 probes targeting 24 AML genes. The dots represent the UMIs of target genes detected by scRaCH-seq (X-axis) and scRNA-seq (Y-axis) for an AML sample.

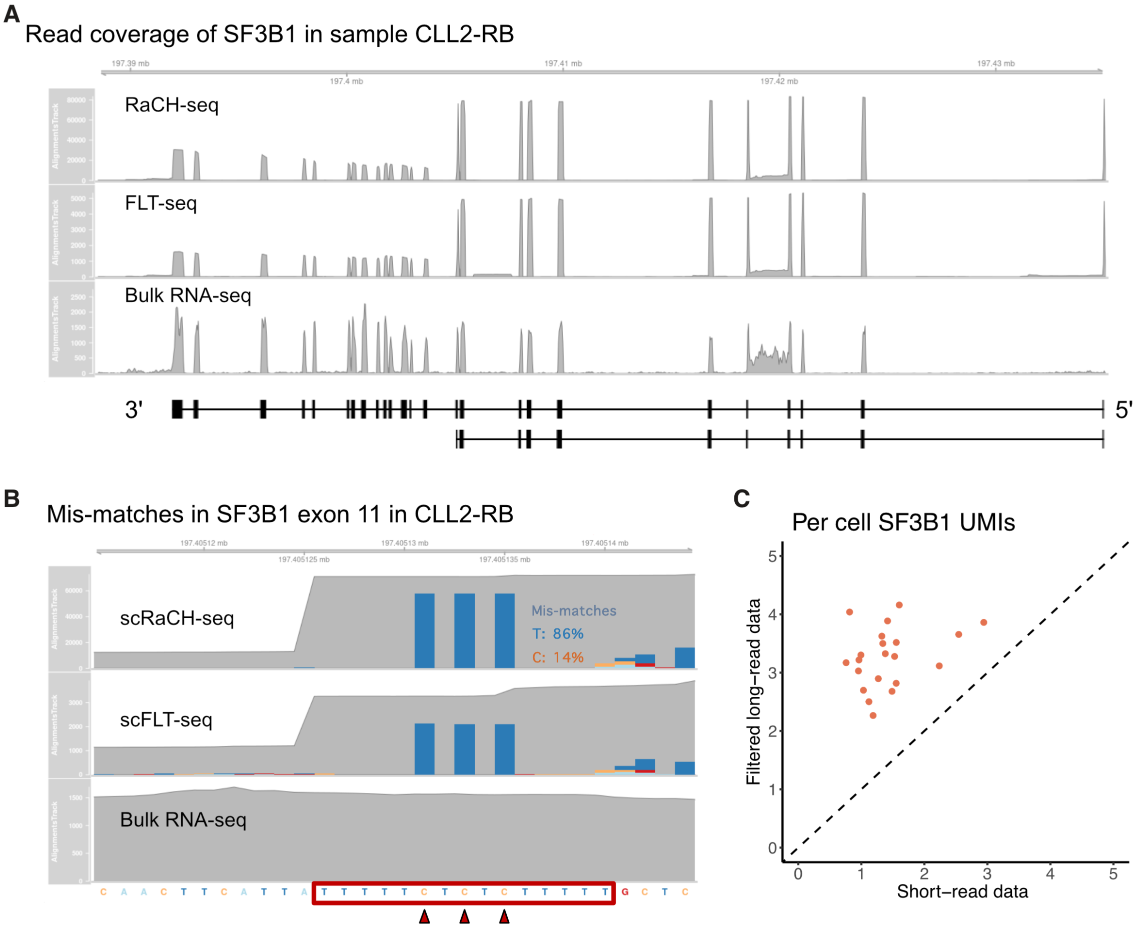


**Supplementary Figure 3. Identification of an SF3B1 artefact introduced by 10x Genomics.**
**A)** Graph showing the SF3B1 read coverage per different sequencing method for sample CLL2-RB.
**B)** Graph showing the mismatches C→T mapped to SF3B1 exon 11 in scRaCH-seq, scFLT and bulk RNA-seq data.
**C)** Dot plot showing the per cell SF3B1 UMIs captured in scRNA-seq (X-axis) and scRaCH-seq (Y-axis) data after artefact removal.

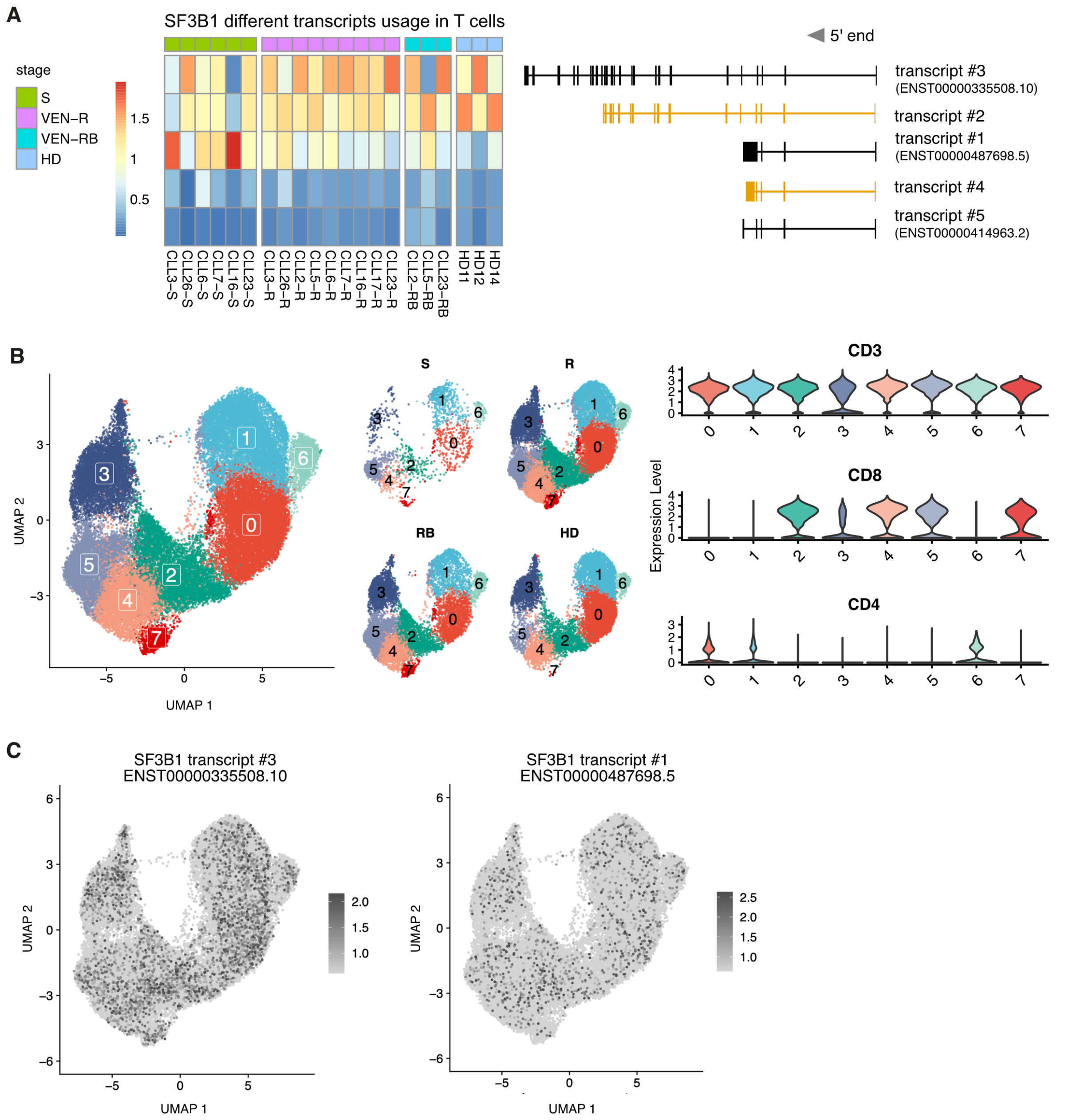


**Supplementary Figure 4 SF3B1 isoform usage by T cells from CLL patients**

**A**) Heatmap showing SF3B1 isoform usage (rows) per sample (columns) in the T cells. The top 5 SF3B1 transcripts are illustrated on the right with novel transcripts highlighted in yellow. Samples are grouped in screening (S; green), venetoclax relapsed (VEN-R/R; pink), venetoclax relapsed and subsequently on BTKi (VEN-RB/RB; blue) and healthy donor (HD; purple).

**B**) UMAP projection of T cells from CLL patients and healthy donors and clustering based on short-read gene expression data. Middle panel showing the UMAP projection of T cells per treatment group. Violin plot showing the expression of the T cell marker (CD3), CD8 T cell marker (CD8) and CD4 T cell marker (CD4) per cluster.

**C**) UMAP projection of SF3B1 transcript #1 and #3 gene-level expression (dark grey) in the T cells.

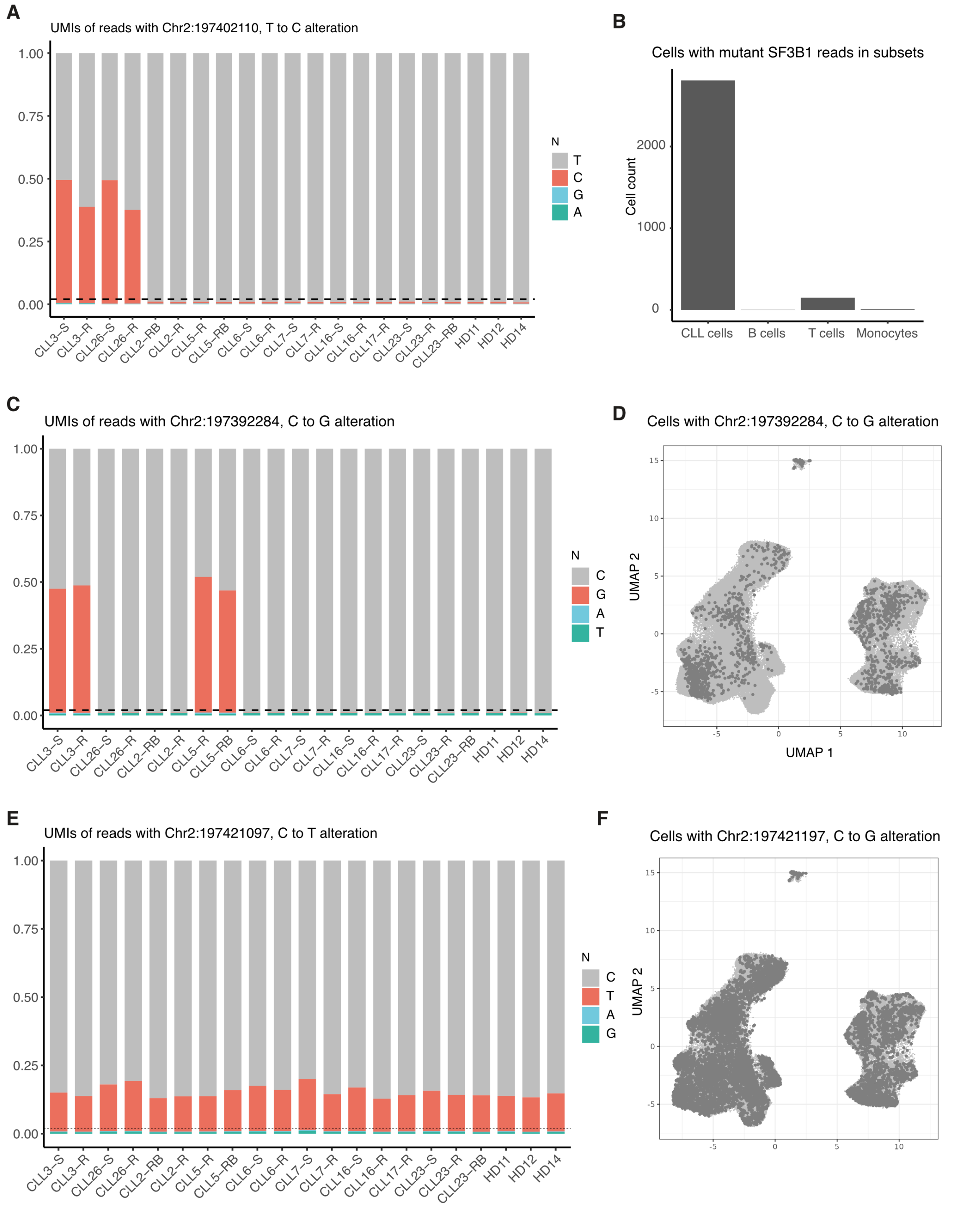


**Supplementary Figure 5 SF3B1 altered transcript expression in CLL and B cells.**

**A**) Bar plot showing the abundance of nucleotide (A, T, C, G) at Chr2:197402110 per sample. The dashed horizontal line indicates the baseline sequencing error (3%) of nanopore sequencing.

**B**) Bar plot showing the quantification of cells with T-to-C alterations at Chr2:197402110 in CLL and non-CLL clusters.

**C**) Bar plot showing the abundance of nucleotide (A, T, C, G) at Chr2:197392284 per sample. The dashed horizontal line indicates the baseline sequencing error (3%) of nanopore sequencing.

**D**) UMAP projection of cells carrying the Chr2:197392284 C-to-G alteration (dark grey)

**E**) Bar plot showing the abundance of nucleotide (A, T, C, G) at Chr2:197421097 per sample. The dashed horizontal line indicates the baseline sequencing error (3%) of nanopore sequencing.

**F**) UMAP projection of cells carrying the Chr2:197421097 C-to-G alteration (dark grey).

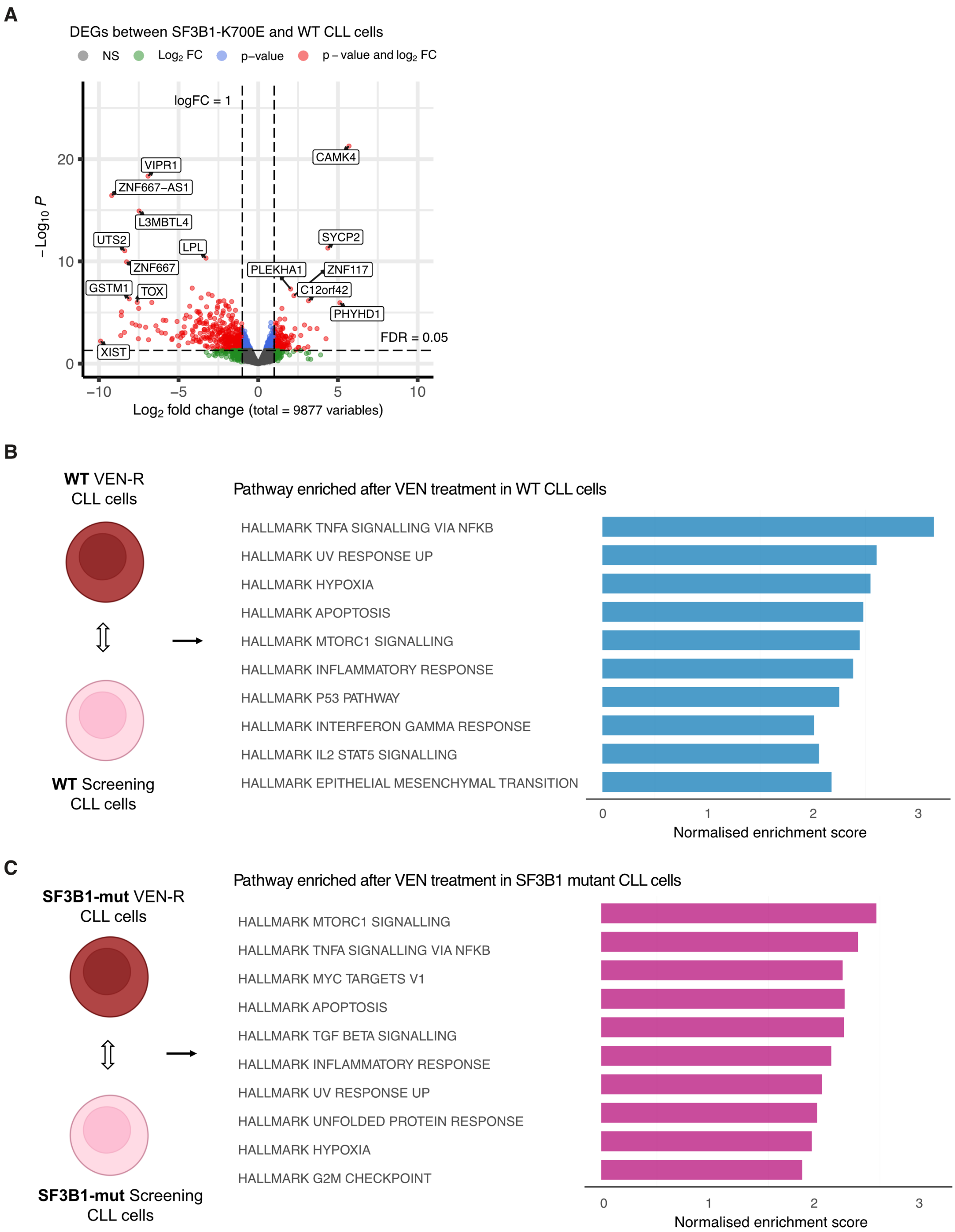


**Supplementary Figure 6 Altered transcriptional expression in SF3B1 K700E mutated cells.**
**A)** Volcano plot showing the differentially expressed genes (DEGs; FDR<0.05) found by pseudo-bulk DE analysis between SF3B1 K700E mutant and wild-type CLL cells.
**B**) Graph showing the enriched pathways in wild-type SF3B1 venetoclax-relapsed CLL cells (blue). CLL cells with >10 wild-type SF3B1 transcripts were used for differential gene expression analysis between the venetoclax-relapsed (VEN-R) group to the screening group. The differentially expressed genes were used as input for gene set enrichment analysis.
**C)** Graph showing the enriched pathways in SF3B1 K700E mutant venetoclax-relapsed CLL cells (pink). CLL cells with >2 mutant SF3B1 transcripts were used for differential gene expression analysis between the venetoclax-relapsed (VEN-R) group to the screening group. The differentially expressed genes were used as input for gene set enrichment analysis.
